# Conserved variation across scales unveils dialectical relationships of micro- and macroevolution

**DOI:** 10.1101/2024.09.02.610914

**Authors:** Keita Saito, Masahito Tsuboi, Yuma Takahashi

## Abstract

Variation enables short-term evolution (microevolution), but its role in long-term evolution (macroevolution) is debated. Here, we analyzed a dataset of *Drosophila* wing variation across six levels of biological organization to demonstrate that microevolutionary variation and macroevolutionary divergence are positively correlated at all levels from variation within an individual to 40 million years of macroevolution. Surprisingly, the strongest relationship was between developmental noise and macroevolutionary divergence—levels thought to be the most distant—whereas the relationship between standing genetic variation and population divergence was modest, despite established theoretical predictions. Our results indicate that the congruence of developmental system with long-term history of fluctuation in adaptive peaks creates dialectical relationships between microevolution and macroevolution.

## Introduction

Since the modern synthesis of evolutionary biology, it has been argued that microevolutionary processes of mutation, selection, genetic drift, and gene flow operating within populations can explain, or at least are consistent with, evolution at long timescales (*1*). Although debates on the role of additional processes operating above the population level remain (*2*, *3*), the extrapolating view of macroevolution profoundly influences our current thinking of evolution (*4*). Arguments in favor of this view rest on evidence from evolutionary genetics. Based on previous studies, the pattern of phenotypic divergence in high-dimensional phenotype space is biased in directions that harbor high amounts of additive genetic variance (*5–7*). Rapidly accumulating evidence indicates that macroevolutionary divergence in various traits and taxa is predictable from standing genetic variation in contemporary populations (*8–11*), as predicted if genetic constraints are relevant for macroevolution. The increasing evidence for this constraint hypothesis supports the extrapolating view of macroevolution, but the mechanisms underlying this inference remain elusive (e.g., “the paradox of predictability,” Tsuboi et al. in press).

The paradox of predictability could be reconciled if variability—the ability of a developmental system to produce variation (*12*)—mirrors the pattern of long-term fluctuations in adaptive peaks. This idea, hereafter referred to as the congruence hypothesis, challenges the established theory of evolutionary genetics by arguing that the standing genetic variation of a population (i.e., evolvability)(*13*) is a consequence, rather than a cause, of macroevolution. Two lines of evidence support this hypothesis. First, the pattern of phenotypic plasticity is consistent with a major axis of phenotypic divergence through computer simulations (*14*), laboratory experiments (*15*), and meta-analyses (*16*). Second, Rohner and Berger (*17*) have recently shown that the subtle responses of development to random and localized environmental fluctuations (i.e., developmental noise)(*18*, *19*) are correlated with the pattern of macroevolutionary divergence. The congruence hypothesis offers a new class of mechanisms regarding the role of non-genetic variations in evolution (*20–23*), which is consistent with the established theory of genetics (*24*, *25*).

In evaluating the constraints and congruence hypotheses, measures of variance at different levels of biological organization are necessary. Here, a multivariate approach was used to estimate the pattern of variation in the wing morphology of *Drosophila* flies at six levels: (1) species divergence, (2) population divergence within a species (*D. simulans*), (3) genetic variation within a population, (4) phenotypic plasticity, (5) mutational variance, and (6) developmental noise. Each variation was summarized as the co/variance matrix and expressed as follows: **R**, wider species divergence; **D**_sp_, narrow species divergence; **D**_pop_, population divergence; **G**, genetic variation; **E**, phenotypic plasticity; **M**, mutational variance; **F**, developmental noise. Using theoretical frameworks of quantitative genetics (*26*, *27*) and statistical physics (*28*) as our guiding principles, we dissect the relationship of variations across these six levels to elucidate the mechanisms underlying the relationship among variability, variation, microevolution, and macroevolution.

## Results and Discussion

### Correlation between mutational variance and fluctuating asymmetry

Two theoretical frameworks have been proposed to elucidate the relationship between variation and divergence. Based on the quantitative genetic model of Lande (*26*), the first framework predicts a positive relationship between the additive genetic co/variance matrix (**G)** and divergence among conspecific populations. If these constraints remain stable for a long period of time and the drift dictates evolution, then the pattern of genetic constraints characterized by **G** scales up to the divergence across species and higher taxa (*1*). The second framework based on the quantitative genetic model of Lynch and Hill (*27*) and the statistical physics model of Kaneko and Furusawa (*28*) indicate that variability is proportional to the pattern of long-term evolutionary divergence among species.

Traditionally, variability has been measured as the subset of phenotypic variance attributable to spontaneous mutation, which is summarized in the mutational co/variance matrix, **M** (*29*). Based on **M**, Lynch and Hill (*27*) proposed that the rate of macroevolution (**R**) is proportional to 2**M**:

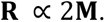

In a related model, Kaneko and Furusawa (*28*) assumed variability as developmental noise, and they proposed that the rate of evolution is proportional to the variance of developmental noise 〈(*δX*)^2^〉. This evolutionary fluctuation–response relationship can be expressed as follows:

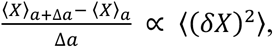

where the variance of the phenotypic trait *X* for a given system is parameterized by *a*, which is assigned as a parameter that specifies the genotype in this study. Both models predict positive correlations between variability and the rate of macroevolution. A technical difference is found when the two variabilities are measured. **M** is estimated from mutation accumulation experiments (*30*), whereas developmental noise can be measured as the random difference between the left- and right sides of laterally symmetric homologs (fluctuating asymmetry, FA) (*31*–*33*). FA represents several favorable attributes as a measurement of variability (*17*), but to operationalize FA in the context of genetics, we need to establish that it is correlated with heritable variability (i.e., **M**).

Thus, we compared **M** in *D. melanogaster*, estimated by Houle and Fierst (*34*) and Houle et al. (*8*), and the co/variance matrix that represents the variation caused by FA (**F**). Two types of **M** were used: spontaneous mutational variance measured under homozygous (**M**_hom_) and heterozygous (**M**_het_) conditions. **F** was estimated using a random mixed-effect model with repeated measurements from one population of *D. simulans*. Linear regression analysis of **M** against **F** revealed a strong positive relationship (Fig. 1; **M**_hom_ on **F**: *R*^2^ = 0.83, *β* = 0.67 ± 0.08; **M**_het_ on **F**: *R*^2^ = 0.95, *β* = 0.73 ± 0.07). Therefore, the two kinds of variabilities considered by Lynch and Hill (*27*) and Kaneko and Furusawa (*28*) may be used to measure the same biological entity. Organismal phenotype is expressed through a developmental system that is dependent on system inputs such as genotype and environmental parameters (*35*, *36*). Given that genotype and environmental parameters are fixed when measuring **M** and **F**, **M** captures the effect of de novo mutation on phenotype during the developmental process, whereas **F** captures the effect of intrinsic developmental perturbation on phenotype during the developmental process. Consequently, both matrices could measure the robustness of phenotype against different sources of perturbations. Based on the framework developed by Lynch and Hill (*27*) and Kaneko and Furusawa (*28*), the robustness of the phenotype during the developmental process could affect the rate of evolution and serve as a primary constraint in the evolutionary process.

**Fig. 1.**
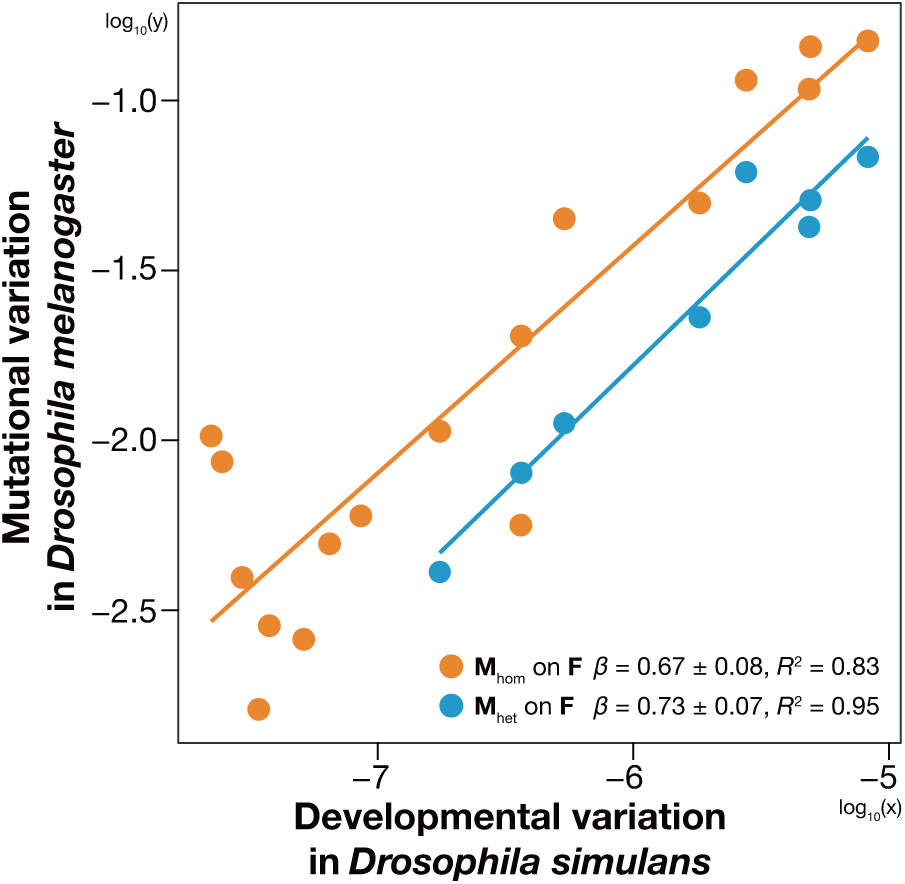
Relationship between two variabilities. Points represent log_10_ (variance in **M** or **F**) along the eigenvectors of **G** in *Drosophila melanogaster*. Key gives log–log regression result, *β* ± s.e. and *R*^2^. Lines indicate single regression lines, and they have slopes equal to *β*. Only upper 17 dimensions of **M**_hom_ and eight dimensions of **M**_het_ were used.

### Variances are positively correlated with divergence at all levels

Next, we examined the validity of the constrains hypothesis (*26*) and the congruence hypothesis (*27*, *28*) by comparing the scaling exponent (i.e., the slope of a log–log regression) and the coefficient of determination (*R*^2^) between divergence and variation or variability. A series of experiments was conducted to estimate the standing variation caused by heritable variation (**G**) and plasticity (**E**). Divergence was evaluated at three levels: divergence among populations of *D. simulans* (**D**_pop_), divergence across 12 *Drosophila* species (**D**_sp_) from an online database (DrosoWing Project) (*37*), and divergence across 112 species of Drosophilidae representing 40 million years of evolution (**R**) from Houle et al. (*8*). If the genetic constraints underpin the relationship between microevolution and macroevolution, then the strongest relationship (i.e., high *R*^2^) and a scaling exponent of 1 are predicted between **G** and **D**_pop_. Conversely, if the congruence hypothesis underlies the micro–macro relationships, then the strongest relationship is predicted between **F** and **R**.

A correlation matrix of the relationships across all pair-wise relationships among variation, variability, and divergence revealed universally positive relationships between the three descriptors of variation (genetic variance, plasticity, and variability) and three levels of divergence (Fig. 2). The correlations are all high (Table 1), which indicates that the variation in the 20-dimensional phenotype space of the *Drosophila* wing morphology is packed into a low-dimensional manifold in a remarkably similar manner, regardless of the causes of variation or the levels of divergence. These similarities are evident in the relative change in the position of each landmark on the wing (Fig. 3). Most landmarks showed a propensity to vary along the lateral axis, particularly landmarks 7, 8, 9, and 10, while the landmarks at the base of the wing exhibited relatively low variation.

**Fig. 2.**
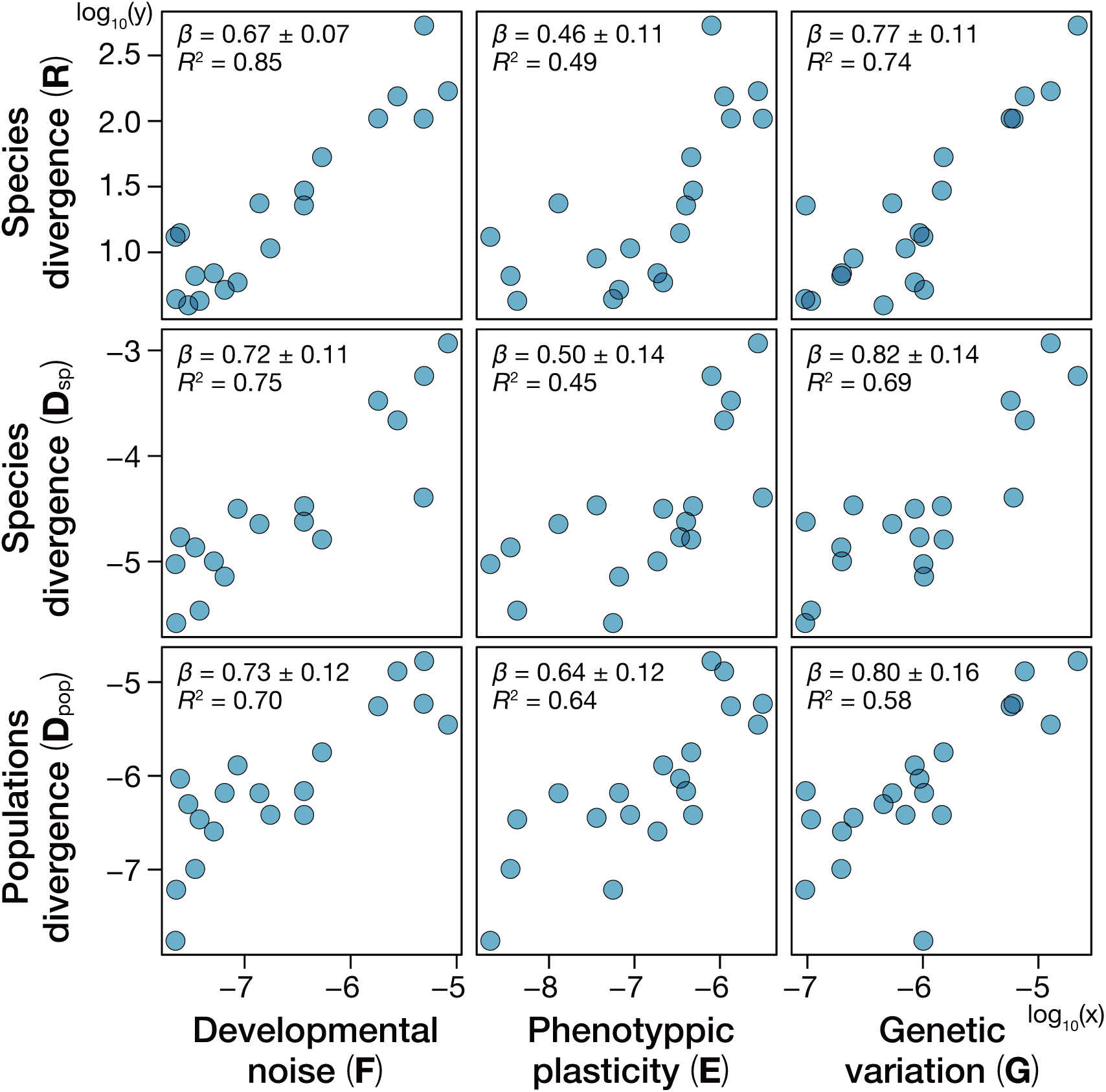
Relationships between variations (G, E, and F) and divergence (D_pop_, D_sp_, and R). Points represent log_10_ (variance in each matrix) along the eigenvectors of **G** in *Drosophila melanogaster*. Key gives log–log regression result, *β* ± s.e. and *R*^2^.

**Fig. 3.**
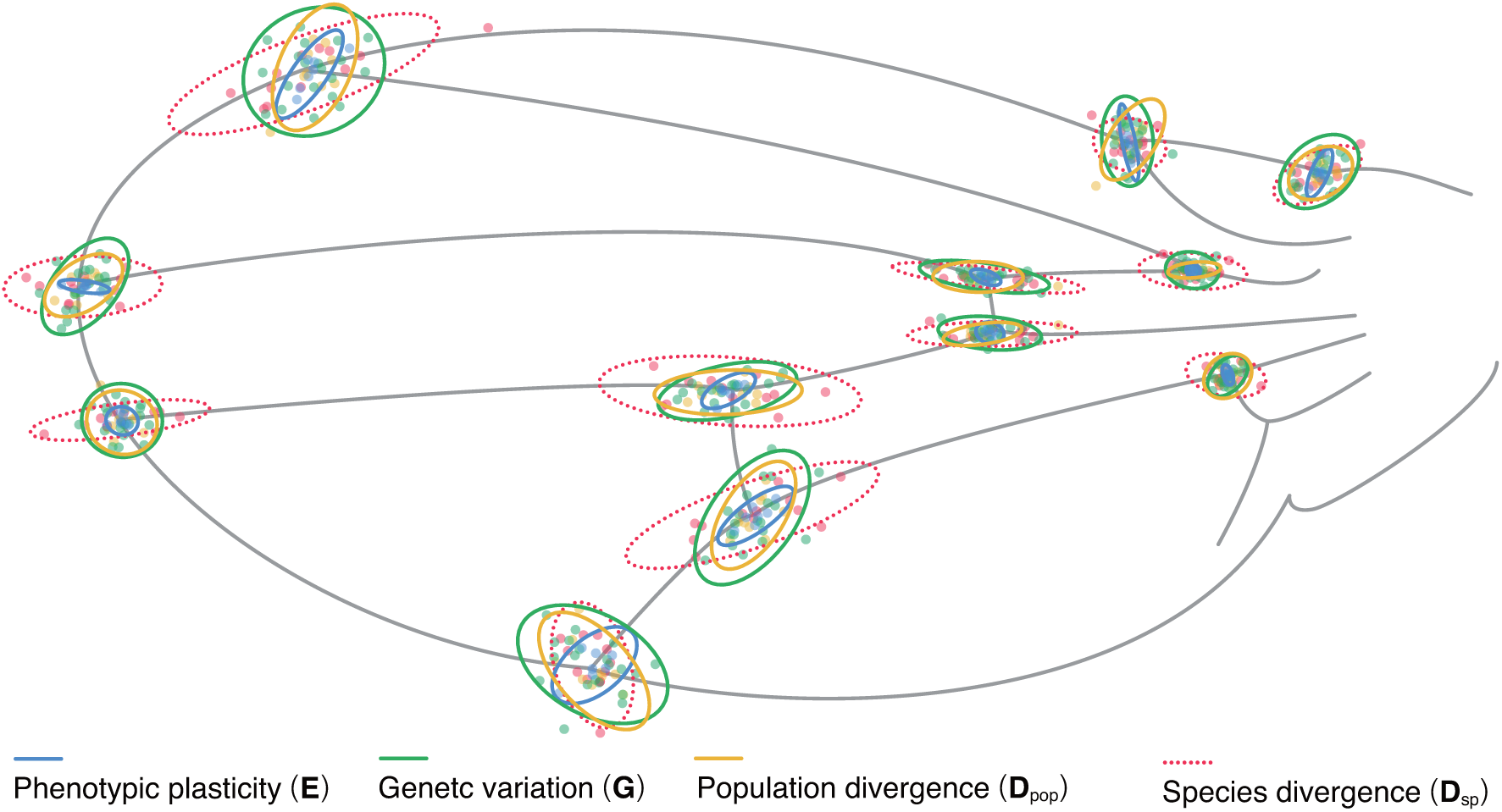
Ellipses representing variation in landmark position. Ellipses are centered, and they represent six SDs (only **D**_sp_: 1.5 SD) for better visibility. Points represent the mean values of species (**D**_sp_), populations (**D**_pop_), genotypes (**G**), and rearing environments (**E**).

**Table 1.** Pearson’s correlation coefficients of pair-wise relationships between log_10_ variances in all six matrices estimated on the basis of MCMCglmm.

| Matrix | 1 | 2 | 3 | 4 | 5 | 6 |
| --- | --- | --- | --- | --- | --- | --- |
| 1. **R** | — |  |  |  |  |  |
| 2. **D<sub>sp</sub>** | 0.87*** | — |  |  |  |  |
| 3. **D<sub>pop</sub>** | 0.78*** | 0.77*** | — |  |  |  |
| 4. **G** | 0.86*** | 0.83*** | 0.76*** | — |  |  |
| 5. **E** | 0.70*** | 0.67** | 0.80*** | 0.66** | — |  |
| 6. **F** | 0.93*** | 0.87*** | 0.84*** | 0.81*** | 0.79*** | — |

Significant codes of *p* values: 0 ‘\*\*\*’ 0.001 ‘\*\*’ 0.01 ‘\*’ 0.05 ‘.’ 1

The comparison of the coefficient of determination (*R*^2^) and slope (*β*) showed that the relationship between **F** and **R** is the strongest (*R*^2^ = 0.85, *β* = 0.67 ± 0.07) among all pair-wise relationships. This result is surprising because these two levels represent the two most distant levels in the biological hierarchy, namely, internal subtle variability of an individual (**F**) and 40 million years of *Drosophila* wing evolution (**R**). Equally surprising is that the relationship between **G** and **D**_pop_ is modest (*R*^2^ = 0.58, *β* = 0.80 ± 0.16), although these two levels are adjacent to each other and have established theoretical and empirical bases to predict a positive relationship (*6*, *26*). These observations support the congruence hypothesis as a leading explanation for the relationship between variation and divergence in phenotypic evolution (*8*–*11*).

### Congruence of variability with macroevolution underlies the pattern of *Drosophila* wing-shaped evolution

The results shown in Fig. 2 are based on the posterior mode of the co/variance matrices estimated at the respective levels (except for **R**). However, considering the difficulty of estimating variation and variability with high precision, some of the results may reflect estimation errors. In examining the impact of the estimation accuracy of each matrix on our inference, the posterior distribution obtained from the Bayesian mixed model was analyzed. The ordinary least-square regressions were used with the co/variance matrices describing each level of divergence presented in Fig. 2 as the response variable and one of the 1,000 posteriors of **G**, **E**, or **F** as the explanatory variable to obtain the distribution of the coefficients of determination (*R*^2^) and the scaling exponent (log–log slope, *β*) of the relationships. After verifying the normality of *R*^2^ and slope distributions of each pair-wise relationship (Table 2), the homoscedasticity and significant differences among variances were examined (Table 3). This analysis (Fig. 4) confirmed that the relationship between **F** and **R** had the highest *R*^2^ value among all pairs, which is significantly higher than the *R*^2^ value between **G** and **D**_pop_.

**Fig. 4.**
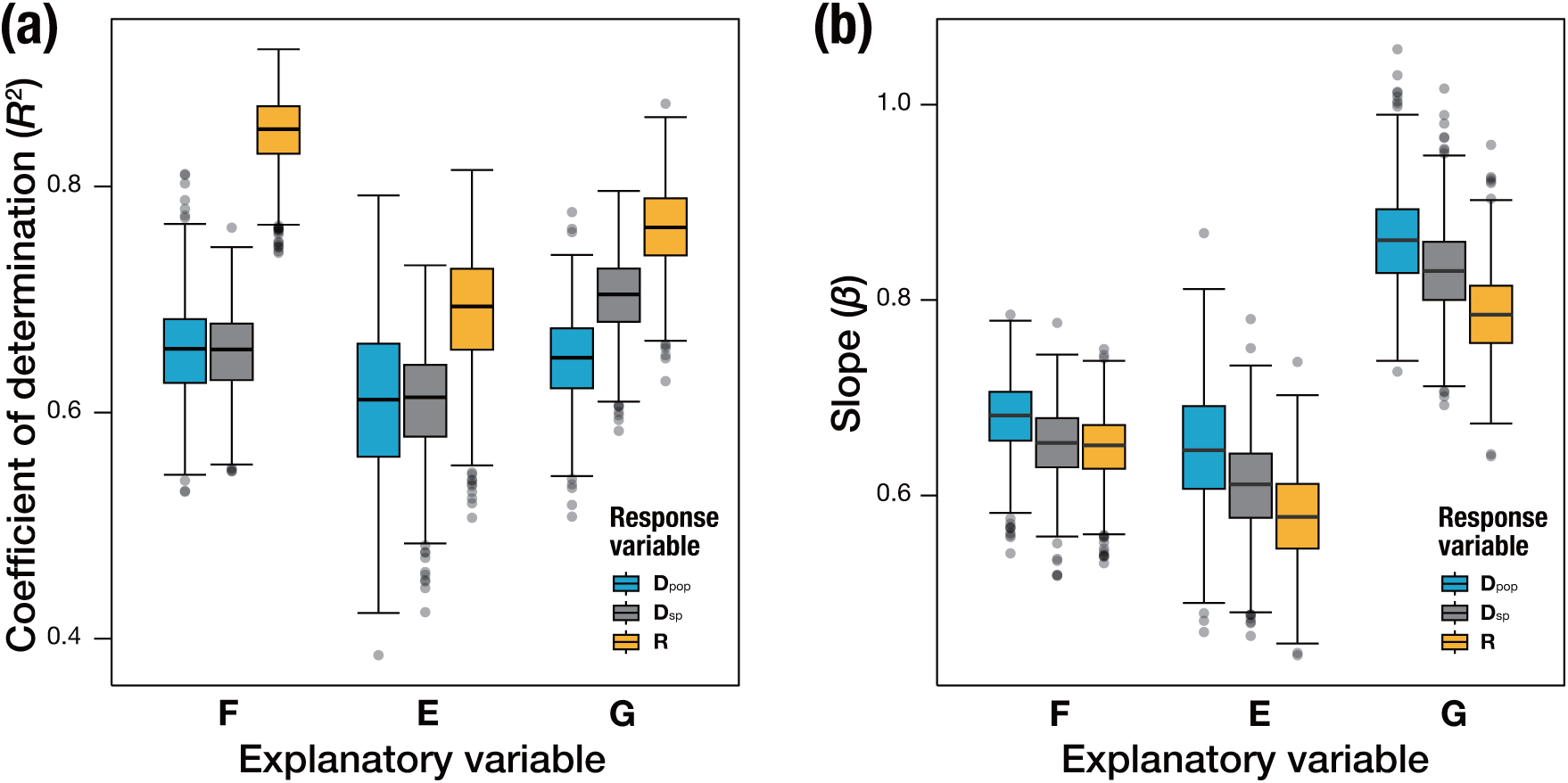
Distributions of *R*^2^ and slope in each regression with variation as explanatory variables and divergence as response variables. Color difference represents the difference in response variables. (a) Distributions of *R*^2^ and (b) distributions of slopes.

**Table 2.** Normality of *R*^2^ and slope variance.

| Response | Explanatory | Normality of $R^2$ | Normality of slopes |
| --- | --- | --- | --- |
| **R** | **G** | $P < 0.05$ | $P < 0.05$ |
| | **E** | $P < 0.001$ | $P = 0.28$ |
| | **F** | $P < 0.001$ | $P < 0.001$ |
| **D<sub>sp</sub>** | **G** | $P < 0.05$ | $P < 0.001$ |
| | **E** | $P < 0.001$ | $P = 0.21$ |
| | **F** | $P = 0.10$ | $P < 0.01$ |
| **D<sub>pop</sub>** | **G** | $P < 0.05$ | $P < 0.01$ |
| | **E** | $P = 0.17$ | $P = 0.38$ |
| | **F** | $P < 0.05$ | $P < 0.01$ |

**Table 3.** Homoscedasticity and the differences among variances.

| $R^2$ | | | | | | |
| --- | --- | --- | --- | --- | --- | --- |
| Response |  |  | Homoscedasticity | T test | U test |  |
| **R** | **G** | **E** | False | $P < 0.001$ | — | |
| | **G** | **F** | False | $P < 0.001$ | — | |
| | **E** | **F** | False | $P < 0.001$ | — | |
| **D<sub>sp</sub>** | **G** | **E** | False | $P < 0.001$ | — | |
| | **G** | **F** | — | — | $P < 0.001$ | |
| | **E** | **F** | — | — | $P < 0.001$ | |
| **D<sub>pop</sub>** | **G** | **E** | — | — | $P < 0.001$ | |
| | **G** | **F** | False | $P < 0.001$ | — | |
| | **E** | **F** | — | — | $P < 0.001$ | |
| Slope |  |  |  |  |  |  |
| **R** | **G** | **E** | — | — | $P < 0.001$ | |
| | **G** | **F** | False | $P < 0.001$ | — | |
| | **E** | **F** | — | — | $P < 0.001$ | |
| **D<sub>sp</sub>** | **G** | **E** | — | — | $P < 0.001$ | |
| | **G** | **F** | False | $P < 0.001$ | — | |
| | **E** | **F** | — | — | $P < 0.001$ | |
| **D<sub>pop</sub>** | **G** | **E** | — | — | $P < 0.001$ | |
| | **G** | **F** | False | $P < 0.001$ | — | |
| | **E** | **F** | — | — | $P < 0.001$ | |

From a purely statistical point of view, **R** and **F** are the most reliably estimated matrices in our dataset, and this clearly has contributed to a tight relationship between these two matrices. **R** is estimated on the basis of 21,138 wings from 117 taxa using a phylogenetic mixed model to extract phylogenetic variances (*8*), and **F** is measured by controlling many sources of variation and instrumental measurement error to extract variances only attributable to local environmental perturbations. By contrast, other matrices (i.e., **D**_sp_, **D**_pop_, **G**, and **E**) are subjected to substantial errors that should have contributed to the low *R*^2^ value. For example, (i) **D**_sp_ is estimated on the basis of the divergence among 12 species; (ii) **G**, **E**, and **D**_pop_ are measured at a certain time from a few representative lines of a single species; (iii) **G** is known as a “broad-sense” **G** that confounds additive with non-additive effects (e.g., epistasis and dominance), and (iv) **E** is at best a poor representation of the inherent plasticity because the environmental conditions that are tolerable by the reared genotypes of *D. simulans* in our experiments are likely a small subset of the range of plasticity available in wild populations of this species (*38*). Given these substantial errors, it is remarkable that all matrices are still unambiguously correlated. Hence, we interpret our results to suggest that all matrices studied here are biologically connected to one another. Rather than placing emphasis on any one of the levels of biological organization as the cause of universal correlations, we favor a dialectical perspective that multiple directions of causes and effects have played important roles in such correlations. Three observations support our view.

First, the tight relationship between **R** and **F** indicates that the macroevolutionary history of fluctuation in adaptive peaks has shaped the developmental system of the *Drosophila* wing. Our causal hypothesis is explicitly that macroevolution determines variability and not *vice versa* because developmental noise is primarily due to local environmental perturbations. Phenotypic variation cannot be inherited without changes in genome. Therefore, the phenotypic variation generated by this process is uninheritable, and does not scale up to variation and divergence at higher levels of biological organization, unlike mutational variance (*5*, *29*).

Second, **F** strongly correlates with **M**, while its correlation with **G** and **E** is positive but modest. Previous studies (*35*, *36*) suggested that development can serve as a mediator that converts different sources of inputs to phenotypic variation. If phenotypic variations caused by different sources are shaped through the same developmental system, then the pattern of phenotypic variation should be similar regardless of whether the variation is genetically or environmentally encoded (*39*). The correlation between **M** and **F** supports this idea and indicates that the propensity of this developmental system to generate variation shapes **G** and **E**. As circumstantial evidence of this, the correlation between **G** and **E** (*r* = 0.66, *P* < 0.01) roughly corresponds to the product of the correlation between **F** and **G** (*r* = 0.81, *P* < 0.001) and **F** and **E** (*r* = 0.79, *P* < 0.001). This would be expected if **F** causes **G** and **E** independently but in a similar manner.

Third, the pattern of microevolution (**D**_pop_) is correlated with **G** with a scaling exponent of 0.80, which is the closest to 1 among all pairs we examined. The evolutionary genetics theory proposes that the directions and rates of microevolution should be determined by **G** (*40–44*) and that the exponent between **D**_pop_ and **G** should be 1 if drift is the dominant evolutionary force. Conversely, as the influence of local adaptations or long-term fluctuations in adaptive peaks strengthens, the scaling exponent of the variance–divergence relationship should increasingly deviate from 1 (*8*, *11*). Therefore, our results are consistent with the established theory of evolutionary genetics and support the idea that genetic constraints dominate the pattern of microevolution.

The following scenarios are proposed to explain our results: (i) macroevolutionary history (**R** and **D**_sp_) shapes the developmental system and determines variability (**F** and **M**); (ii) the developmental system shapes the standing variation (**G** and **E**) under the influence of local adaptation and drift; (iii) **G** determines the pattern of microevolution (**D**_pop_). Based on this hypothesis, macroevolution is often predictable (*8*, *9*, *11*, *17*) because the long-term pattern of fluctuation in adaptive peaks molds variability. Our hypothesis also explains why microevolution is not always predictable from **G** and contemporary selection (*45*, *46*) because neither microevolution, **G**, nor selection are direct descriptors of variability and long-term peak movements.

## Conclusion

Our data supports the congruence mechanism to explain the correlation between macroevolution and the standing variation within a population. Rather than seeking a single explanation, our perspective advocates for a pluralistic view that acknowledges the multifaceted link between microevolution and macroevolution, encompassing multiple evolutionary processes and mechanisms operating at various levels and directions. Although our hypothesis that macroevolution molds variability may appear radical, it is consistent with classic theories of evolutionary genetics (*5*, *6*, *13*, *26*) and evidence supporting our view is widely found in the literature of non-genetic inheritance (*20*–*23*). Our findings contribute to the development of a unified theory of evolution applicable to all time scales.

## Methods

### Sampling and established isofemale line

*Drosophila simulans* were sampled from different wild populations in Japan, and isofemale lines were established (Table 4). Each isofemale line was repeatedly inbred over several generations to reduce genetic variation within an isofemale line and to remove environmental and maternal effects. Each isofemale line was maintained in standard media on the basis of the method of Fitzpatrick et al. (*47*) (500 mL of H_2_O, 50 g of sucrose, 50 g of dry yeast, 6.5 g of agar, 5.36 g of KNaC_4_H_6_.4H_2_O, 0.5 g of KH_2_PO_4_, 0.25 g of NaCl, 0.25 g of MgCl_2_, 0.25 g of CaCl_2_, and 0.35 g of Fe_2_(SO_4_)·6.9H_2_O) in 170-mL bottles under a 12 light:12 dark cycle at 25°C. In addition, *D. lutescens* were sampled from a single wild population in Japan (the campus of Chiba University: 35° 62′ 79′′ N, 140° 10′ 31′′ E), and isofemale lines were established in the same manner of *D. simulans*.

**Table 4.** Sampling points where we chaptered *D. simulans* and the number of isofemale lines used in this study.

| Sampling city | Isofemale lines | Latitude | Longitude |
| --- | --- | --- | --- |
| Chiba (Chiba University) | 33 | 35° 62' 79" N | 140° 10' 31" E |
| Chiba (Showa Forest) | 6 | 35° 52' 03" N | 140° 28' 19" E |
| Ichigaya | 1 | 35° 68' 85" N | 139° 73' 09" E |
| Kisarazu | 4 | 35° 32' 97" N | 139° 99' 08" E |
| Kochi | 5 | 33° 54' 91" N | 133° 48' 68" E |
| Nagoya | 3 | 35° 15' 40" N | 136° 96' 83" E |
| Osaka | 7 | 34° 59' 58" N | 135° 50' 97" E |
| Sendai | 5 | 38° 26' 10" N | 140° 84' 95" E |
| Tateyama | 2 | 34° 98' 15" N | 139° 85' 56" E |
| Tsukuba | 4 | 36° 05' 22" N | 140° 13' 06" E |
| Tsuno | 1 | 32° 28' 80" N | 131° 46' 51" E |

### Rearing experiments

In quantifying the phenotypic plasticity, previously published data were used (*38*). Larvae of individual *D. simulans* were reared from the egg stage through hatching and metamorphosis to adults under seven combinations of three environmental factors. These combinations consisted of three nutrient conditions (high, intermediate, or low), three light–dark cycle conditions (10L:14D, 12L12D, or 14L:10D), and three temperature conditions (20°C, 23°C, or 26°C). Refer to Saito et al. (*38*) for further details of this experiment.

### Wing collection and datasets

After establishing the isofemale lines, adult females were collected from isofemale lines of *D. simulans* and *D. lutescens*. The left and right wings were separated from their bodies and then directly placed on the glass slide. To flatten the wings, a cover glass was placed on the wings and glued to the glass slide (i.e., dry mount). The wings were imaged using the CMOS camera (Leica MC190 HD, 10 million pixels) of the stereoscopic fluorescence microscope (Leica M165 FC) under the condition in which the wings were lighting up by the torres stand under the glass slide. In addition, wing photos were collected from previous studies. First, as mentioned previously, the datasets of Saito et al. (*38*) were used to estimate **E**. Second, wing photos of 13 *Drosophila* species were collected from DrosoWing Project (*37*) to estimate species divergence. These three wing photo datasets were combined and used for the following analysis. The total number of wing photos was 5,569 (Table 5), and all wings were obtained only from female. In estimating the narrow range of species divergence matrix, wing photos of 14 *Drosophila* species were used. On the contrary, for population divergence, genetic variation, variation caused by phenotypic plasticity, and developmental noise, only *D. simulans* wing photos were used.

**Table 5.**
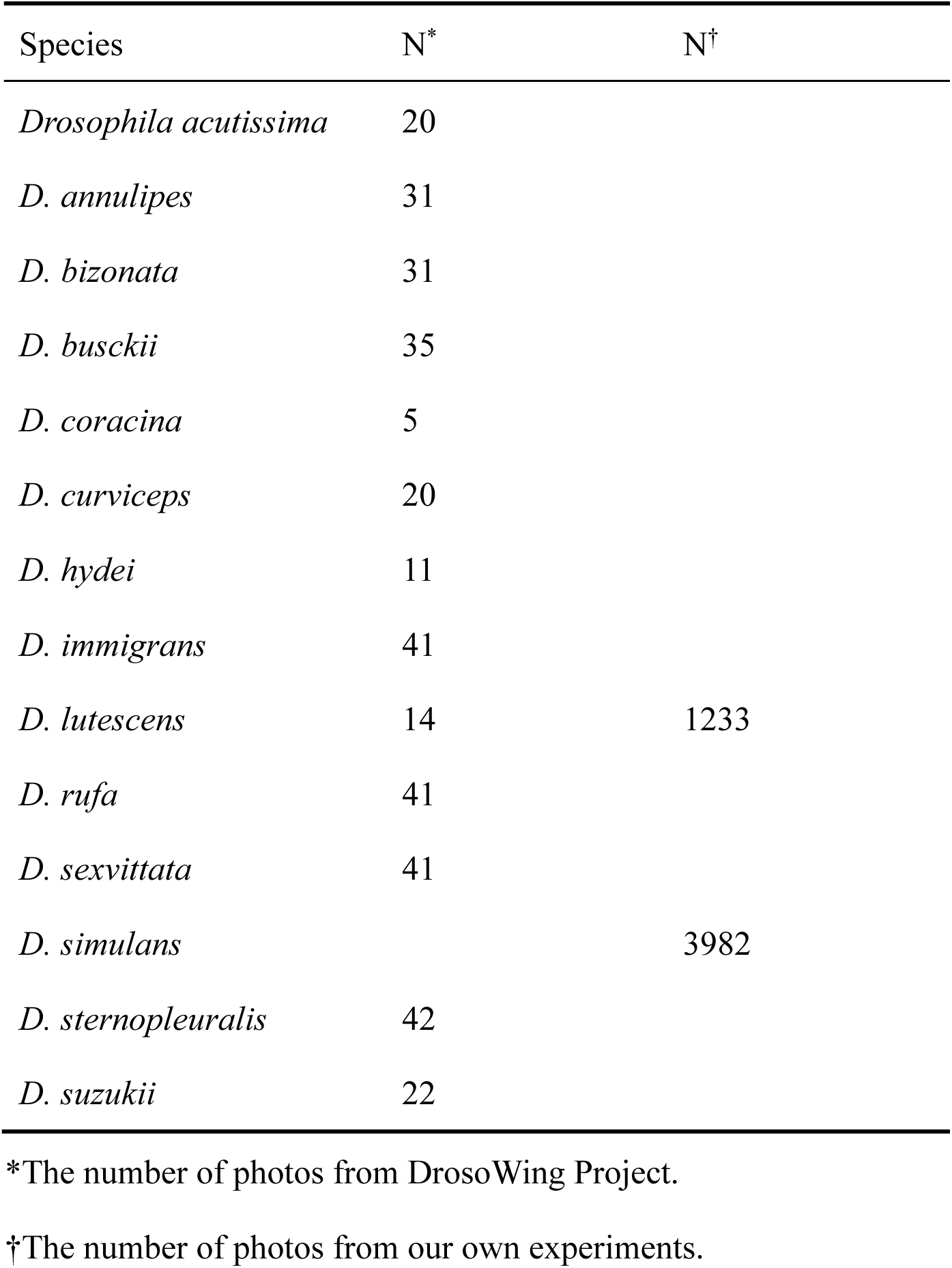
Species name and the number of wing photos.

| Species | N* | N† |
| --- | --- | --- |
| *Drosophila acutissima* | 20 |  |
| *D. annulipes* | 31 |  |
| *D. bizonata* | 31 |  |
| *D. busckii* | 35 |  |
| *D. coracina* | 5 |  |
| *D. curviceps* | 20 |  |
| *D. hydei* | 11 |  |
| *D. immigrans* | 41 |  |
| *D. lutescens* | 14 | 1233 |
| *D. rufa* | 41 |  |
| *D. sexvittata* | 41 |  |
| *D. simulans* |  | 3982 |
| *D. sternopleuralis* | 42 |  |
| *D. suzukii* | 22 |  |

\*The number of photos from DrosoWing Project.
†The number of photos from our own experiments.

### Wing measurements and analyses of wing shapes

In the present study, the *x* and *y* coordinates of 12 landmark vein intersections were measured on the basis of Houle et al. (*8*) from collected photos, and a semi-automated procedure was used to acquire these coordinates. First, the machine-learning program, “*ml-morph*” (*48*), was used to obtain the *x* and *y* coordinates of landmarks automatically. Next, considering that the *x* and *y* coordinates of 12 landmarks extracted from “*ml-morph*” were rarely incorrect, these *x* and *y* coordinates were manually corrected by using “*imglab*” of “*Dlib*.” We also used trained *ml-morph*, which was built in Saito et al. (*38*). This procedure was performed to times to prevent the effect of measurement error. Finally, the *x* and *y* coordinates were geometrically aligned to eliminate the variation caused by wing size, wing direction, and the magnification of the camera when shooting using Generalized Procrustes Analysis (GPA), which translated original *x* and *y* coordinates data to a common coordinate system by keeping constant variation in their position, size, and direction, and were included in the “*geomorph*” package in *R* (*49*). These standardized coordinates were used to estimate each co/variance matrix.

### Estimating variance matrices by MCMCglmm

A Bayesian mixed-effect model implemented in the MCMCglmm package version 2.35 was used to estimate the co/variance matrices at five levels: **D**_sp_, **D**_pop_, **G**, **E**, and **F**. In all models except for the model estimating **F**, standardized *x* and *y* coordinates averaged across four measurements in each individual (i.e., repeated measurements of left and right wing) were analyzed as the multivariate response variable of the model, and a flat uninformative prior was used. Four models were run, each with the random effects of species, population, treatment, or ID of isofemale lines to estimate **D**_sp_, **D**_pop_, **E**, and **G**, respectively. In a model estimating **F**, all measurements (i.e., four measurements per individual) were used as the multivariate response variable in which a dummy variable that index individual ID and side (left or right) serves as the random effect. We did not include side as the fixed effect to estimate directional asymmetry in this model because preliminary analysis showed no effect of side (results not shown). The resultant variance component associated with this dummy variable of the model was used to evaluate FA accounting for instrumental measurement error. As a prior of this model, a matrix was used, whose diagonals are the empirically estimated **F** value reported in Saito et al. (*38*) and off-diagonals are zeros. All models were run for 750,000 iterations with burning and thinning intervals of 250,000 and 5,000, respectively, to yield 1,000 posterior samples. Model convergence was assessed by evaluating the posterior, and chain mixing was tested formally in accordance with the Heidelberg criteria (*50*) where all models passed the test.

### Estimated matrices

For a wider range of species divergence, the rate of macroevolution matrix was used and summarized in **R**. This matrix was estimated in accordance with the method of Houle et al. (*8*). In addition, **M** (**M**_hom_ and **M**_het_) in *D. melanogaster* was used and estimated in accordance with the methods of Houle and Fierst (*34*) and Houle et al. (*8*) to investigate the subset phenotypic variance attributable to spontaneous mutation.

### Comparison

Each co/variance matrix without **R** and **M** was the median of the posterior distribution of the co/variance matrix obtained by each model. In comparing each co/variance matrix, each matrix was projected to the same wing morphospace of the additive genetic co/variance matrix (**G**) of *D. melanogaster* presented in Houle et al. (*8*). Each projected matrix was composed of 24 traits (12 landmarks of *x* and *y* coordinates), but performing GPA reduces the number of dimensions by 4. Therefore, the upper 20 dimensions were used to compare each matrix. On the contrary, **M**_hom_ and **M**_het_ were less than the full rank. Thus, only upper 17 dimensions about **M**_hom_ or eight dimensions about **M**_het_ were used in accordance with the method of Houle et al. (*8*) when we compared **F** and **M**. In examining the relationship among each matrix, single regression was conducted, and the coefficient of determination (R^2^) and the slope score (*β*) were calculated. Each analysis was based on Houle et al. (*8*) and Rohner and Berger (*17*).

## Author contributions

Conceptualization: All authors. Methodology: All authors. Investigation: K.S. Visualization: K.S. and M.T. Funding acquisition: Y.T. Project administration: All authors. Supervision: M.T. and Y.T. Writing – original draft: K.S. Writing – review & editing: All authors.

## Data and materials availability

All raw data and R code used in this study have been uploaded to Figshare, 10.6084/m9.figshare.26861356.

## References

1. B. Charlesworth, R. Lande, M. Slatkin, A Neo-Darwinian Commentary on Macroevolution. Evolution 36, 474–498 (1982).

2. D. Jablonski, Approaches to macroevolution: 2. sorting of variation, some overarching issues, and general conclusions. Evol Biol 44, 451–475 (2017).

3. M. Kearney, B. S. Lieberman, L. C. Strotz, Tangled banks, braided rivers, and complex hierarchies: beyond microevolution and macroevolution. Journal of Evolutionary Biology, voae065 (2024).

4. D. J. Futuyma, M. Kirkpatrick, Evolution (Sinauer Associates, Inc., Publishers, Sunderland, Massachusetts, Fourth edition., 2017).

5. S. J. Arnold, Constraints on phenotypic evolution. The American Naturalist 140, 85–107 (1992).

6. D. Schluter, Adaptive radiation along genetic lines of least resistance. Evolution 50, 1766–1774 (1996).

7. B. Walsh, M. W. Blows, Abundant genetic variation + strong selection = multivariate genetic constraints: a geometric view of adaptation. Annual Review of Ecology, Evolution, and Systematics 40, 41–59 (2009).

8. D. Houle, G. H. Bolstad, K. van der Linde, T. F. Hansen, Mutation predicts 40 million years of fly wing evolution. Nature 548, 447–450 (2017).

9. J. W. McGlothlin, M. E. Kobiela, H. V. Wright, D. L. Mahler, J. J. Kolbe, J. B. Losos, E. D. Brodie, Adaptive radiation along a deeply conserved genetic line of least resistance in *Anolis* lizards. Evolution Letters 2, 310–322 (2018).

10. Ø. H. Opedal, W. S. Armbruster, T. F. Hansen, A. Holstad, C. Pélabon, S. Andersson, D. R. Campbell, C. M. Caruso, L. F. Delph, C. G. Eckert, Å. Lankinen, G. M. Walter, J. Ågren, G. H. Bolstad, Evolvability and trait function predict phenotypic divergence of plant populations. Proceedings of the National Academy of Sciences 120, e2203228120 (2023).

11. A. Holstad, K. L. Voje, Ø. H. Opedal, G. H. Bolstad, S. Bourg, T. F. Hansen, C. Pélabon, Evolvability predicts macroevolution under fluctuating selection. Science 384, 688–693 (2024).

12. G. P. Wagner, L. Altenberg, Complex adaptations and the evolution of evolvability. Evolution 50, 967–976 (1996).

13. T. F. Hansen, D. Houle, M. Pavličev, C. Pélabon, Eds., Evolvability: A Unifying Concept in Evolutionary Biology? (The MIT Press, 2023; https://direct.mit.edu/books/book/5602/EvolvabilityA-Unifying-Concept-in-Evolutionary).

14. C. Furusawa, K. Kaneko, Formation of dominant mode by evolution in biological systems. Phys. Rev. E 97, 042410 (2018).

15. C. Furusawa, K. Kaneko, Global relationships in fluctuation and response in adaptive evolution. Journal of The Royal Society Interface 12, 20150482 (2015).

16. D. W. A. Noble, R. Radersma, T. Uller, Plastic responses to novel environments are biased towards phenotype dimensions with high additive genetic variation. Proceedings of the National Academy of Sciences 116, 13452–13461 (2019).

17. P. T. Rohner, D. Berger, Developmental bias predicts 60 million years of wing shape evolution. Proceedings of the National Academy of Sciences 120, e2211210120 (2023).

18. J. L. Spudich, D. E. Koshland, Non-genetic individuality: chance in the single cell. Nature 262, 467–471 (1976).

19. T. Uller, A. P. Moczek, R. A. Watson, P. M. Brakefield, K. N. Laland, Developmental bias and evolution: a regulatory network perspective. Genetics 209, 949–966 (2018).

20. T. D. Price, A. Qvarnström, D. E. Irwin, The role of phenotypic plasticity in driving genetic evolution. Proceedings of the Royal Society B: Biological Sciences 270, 1433–1440 (2003).

21. C. K. Ghalambor, J. K. McKAY, S. P. Carroll, D. N. Reznick, Adaptive versus non adaptive phenotypic plasticity and the potential for contemporary adaptation in new environments. Functional Ecology 21, 394–407 (2007).

22. E. Danchin, Avatars of information: towards an inclusive evolutionary synthesis. Trends in Ecology & Evolution 28, 351–358 (2013).

23. J. Draghi, Developmental noise and ecological opportunity across space can release constraints on the evolution of plasticity. Evolution & Development 22, 35–46 (2020).

24. G. P. Wagner, G. Booth, H. Bagheri-Chaichian, A Population genetic theory of canalization. Evolution 51, 329–347 (1997).

25. J. A. Draghi, M. C. Whitlock, Phenotypic plasticity facilitates mutational Variance, genetic variance, and evolvability along the major axis of environmental variation. Evolution 66, 2891–2902 (2012).

26. R. Lande, Quantitative genetic analysis of multivariate evolution, applied to brain: body size allometry. Evolution 33, 402–416 (1979).

27. M. Lynch, W. G. Hill, Phenotypic evolution by neutral mutation. Evolution 40, 915–935 (1986).

28. K. Kaneko, C. Furusawa, Macroscopic theory for evolving biological systems akin to thermodynamics. Annual Review of Biophysics 47, 273–290 (2018).

29. R. Lande, The genetic covariance between characters maintained by pleiotropic mutations. Genetics 94, 203–215 (1980).

30. D. L. Halligan, P. D. Keightley, Spontaneous mutation accumulation studies in evolutionary genetics. Annual Review of Ecology, Evolution, and Systematics 40, 151–172 (2009).

31. L. Van Valen, A study of fluctuating asymmetry. Evolution 16, 125–142 (1962).

32. Gangestad, Thornhill, Individual differences in developmental precision and fluctuating asymmetry: a model and its implications. Journal of Evolutionary Biology 12, 402–416 (1999).

33. P. T. Rohner, Y. Hu, A. P. Moczek, Developmental bias in the evolution and plasticity of beetle horn shape. Proceedings of the Royal Society B: Biological Sciences 289, 20221441 (2022).

34. D. Houle, J. Fierst, Properties of spontaneous mutational variance and covariance for wing size and shape in Drosophila melanogaster. Evolution 67, 1116–1130 (2013).

35. M.-A. Félix, M. Barkoulas, Pervasive robustness in biological systems. Nat Rev Genet 16, 483–496 (2015).

36. C. P. Klingenberg, Phenotypic plasticity, developmental instability, and robustness: the concepts and how they are connected. Frontiers in Ecology and Evolution 7, 1–15 (2019).

37. S. Y. M. Loh, Y. Ogawa, S. Kawana, K. Tamura, H. K. Lee, Semi-automated quantitative *Drosophila* wings measurements. BMC Bioinformatics 18, 319 (2017).

38. K. Saito, M. Tsuboi, Y. Takahashi, Developmental noise and phenotypic plasticity are correlated in *Drosophila simulans*. Evolution Letters, qrad069 (2024).

39. D. Houle, Comparing evolvability and variability of quantitative traits. Genetics 130, 195–204 (1992).

40. M. W. Blows, M. Higgie, A. E. D. E. L. Promislow, Genetic constraints on the evolution of mate recognition under natural selection. The American Naturalist 161, 240–253 (2003).

41. G. Marroig, J. M. Cheverud, Size as a line of least evolutionary resistance: diet and adaptive morphological radiation in New World monkeys. Evolution 59, 1128–1142 (2005).

42. T. F. Hansen, D. Houle, Measuring and comparing evolvability and constraint in multivariate characters. Journal of Evolutionary Biology 21, 1201–1219 (2008).

43. S. F. Chenoweth, H. D. Rundle, M. W. Blows, The contribution of selection and genetic constraints to phenotypic divergence. The American Naturalist 175, 186–196 (2010).

44. R. I. Colautti, S. C. H. Barrett, Population divergence along lines of genetic variance and covariance in the invasive plant *Lythrum salicaria* in eastern North America. Evolution 65, 2514–2529 (2011).

45. B. Pujol, S. Blanchet, A. Charmantier, E. Danchin, B. Facon, P. Marrot, F. Roux, I. Scotti, C. Teplitsky, C. E. Thomson, I. Winney, The missing response to selection in the wild. Trends in Ecology & Evolution 33, 337–346 (2018).

46. M. C. Urban, J. Swaegers, R. Stoks, R. R. Snook, S. P. Otto, D. W. A. Noble, M. Moiron, M. H. Hällfors, M. Gómez-Llano, S. Fior, J. Cote, A. Charmantier, E. Bestion, D. Berger, J. Baur, J. M. Alexander, M. Saastamoinen, A. H. Edelsparre, C. Teplitsky, When and how can we predict adaptive responses to climate change? Evolution Letters 8, 172–187 (2024).

47. M. J. Fitzpatrick, E. Feder, L. Rowe, M. B. Sokolowski, Maintaining a behaviour polymorphism by frequency-dependent selection on a single gene. Nature 447, 210–212 (2007).

48. A. Porto, K. L. Voje, ML-morph: a fast, accurate and general approach for automated detection and landmarking of biological structures in images. Methods in Ecology and Evolution 11, 500–512 (2020).

49. D. C. Adams, E. Otárola-Castillo, geomorph: an R package for the collection and analysis of geometric morphometric shape data. Methods in Ecology and Evolution 4, 393–399 (2013).

50. P. Heidelberger, P. D. Welch, Simulation run length control in the presence of an initial transient. Operations Research 31, 1109–1144 (1983).

